# Computational exploration of bio-remediation solution for mixed plastic waste

**DOI:** 10.1101/2022.03.20.485065

**Authors:** Sunny, Ankita Maurya, Mohit Kumar Vats, Sunil Kumar Khare, Kinshuk Raj Srivastava

## Abstract

The plastic materials are recalcitrant in the open environment, surviving longer without complete remediation. The current disposal methods of used plastic material are not efficient; consequently, plastic wastes are infiltrating the natural resources of the biosphere. A sustaining solution for plastic waste is either recycling or making it part of the earth’s biogeochemical cycle. We have collected, manually mined, and analyzed the previous reports on plastic biodegradation. Our results demonstrate that the biodegradation pattern of plastics follows the chemical classification of plastic types. Based on clustering analysis, the distant plastic types are grouped into two broad categories of plastic types, C-C (non-hydrolyzable) and C-X (hydrolyzable). The genus enrichment analysis suggests that Pseudomonas and Bacillus from bacteria and Aspergillus and Penicillium from fungal are potential genera for bioremediation of mixed plastic waste. Overall results have pointed towards a possible solution of mixed plastic waste either in a circular economy or open remediation. The meta-analysis of the reports revealed a historical inclination of biodegradation studies towards C-X type of plastic; however, the C-C class is dominated in overall plastic production. An interactive web portal of reports is hosted at plasticbiodegradation.com for easy access by other researchers for future studies

## Introduction

Plastic is a collective term used for multiple polymeric materials known to be lightweight, durable, and simultaneously can own bespoke shapes. A polymeric material is an extended repetition of the characterized monomeric unit synthesis from nature or chemical origins. Besides having advanced materialistic properties, they also provide an economical and real-time supply edge over alternatives such as glass, steel, and bio-based fibers. Naphtha and natural gas are the leading suppliers of plastic monomers, and it accounts for 6% of total annual crude oil productions (Geyer *et al*., 2017). There are around 50 main plastic types; however, their annual production is not uniform but skewed towards a few popular plastic types (Ramkumar *et al*., 2021). The top 6 popular plastic types, polyethylene (PE), polypropylene (PP), poly(vinyl chloride) (PVC), polystyrene (PS), poly(ethylene terephthalate) (PET), and polyurethane (PU), account for more than three-quarters of total plastic production (Hauer *et al*. 2020; Danso *et al*., 2019). The overall layout of the polymerization process can yield a linear or branched polymer with a varying grade of crystallinity which dictates the usability profile of a polymer (Debuissy *et al*., 2018). Plastics available in the market are not homogenous but contain additives, dyes, plasticizers, antioxidants, and antibiotics to achieve the tailor’s properties (Wiesinger *et al*., 2021; Zimmermann *et al*., 2019). Based on the combination of atoms in the main backbone, we can divide plastic into C-C or C-X groups, where X is any heteroatom other than carbon (Inderthal *et al*., 2021); Gewert *et al*., 2015). Within the C-C group are the vinyl polymers, which are highly resistant to any form of degradation due to the inert nature of their backbone. The C-X class has multiple subtypes based on the nature of the bond between carbon and heteroatom, such as ether, amide, ester. Figure1 shows the hierarchical classification of plastic types with examples.

The annual production of plastic has surged from 1.5 million metric tons in 1950 to 368 million metric tons in 2015, indicating the global economic dominance of plastic in a short period. (Geyer et al., 2017). The success of plastic as a wonder material paved the path of modern civilization such that it has become a significant part of every aspect of our lives, from personal to the global economy. The monopoly of plastic material leads to prolonged negligence of the end fate of plastic products, which has marked an Anthropocene Epoch in earth’s history, where we can observe a ubiquitous presence of plastic litter from the deepest part of the ocean surface to the high altitude region of the mountains (Geyer *et al*., 2017, Napper *et al*., 2020; Raddadi & Fava, 2019). The current state of plastic waste management is highly underutilized (Vollmer *et al*., 2020). The typical approach of reducing, redesigning, reusing, and recycling is not working because this is not making any dent in the futuristic production of plastic waste material (Maurya *et al*., 2020; Wei *et al*., 2020). The recent estimates point towards a situation of less than 20% overall recycling of plastic waste (Geyer *et al*., 2017). These recycling efforts do not resupply the same plastic types used in the process but instead become the source of downgraded plastics; meanwhile, the supply of recycled plastic types is continuously coming from virgin plastic synthesis. The single-use plastic contributes around 50% of the overall plastic production and frequently ends up in the open environment due to poor incentives to recycle and irresponsible consumer behaviour. It has become the most-talked-about category of plastic in terms of regulation and management (Xanthos & Walker, 2017; Elliott *et al*., 2020). The current global supply chain has made the scenario even more complex as the producer and end-user often fall in different areas of the globe, so any legal options to pass the liability on producer has a thin margin for emerging (Lebreton & Andrady, 2019).

Plastic waste has a short history in the open environment to get a proper (functional) natural biochemical cycle in the earth’s ecosystem. It is an unparalleled catastrophe of xenobiotics faced by living organisms (Amaral-Zettler *et al*., 2020). The *microbial scavengers* play an essential role in all biogeochemical cycles, and they are robust to react to their ecological niche by evolving their molecular machinery (Fenner *et al*., 2021; Lithner *et al*., 2011,). The plastic waste of size less than 5mm, called microplastic, is making the situation even worse as there is no physical way to stop them from contaminating every fabric of the earth’s environment (Kooi *et al*., 2021; Mitrano & Wohlleben, 2020; Yuan *et al*., 2020). Due to the continued availability of plastic in the open environment, weathering causes fragmentation of plastic material into microplastic, which exacerbates the problem (Verla *et al*., 2019) and have become a hotspot for antibiotic-resistant microorganisms to form biofilms (Kirstein *et al*., 2018; Dussud *et al*., 2018; Jacquin *et al*., 2019), revealing a wide range of antibiotic resistance genes of pathogenic bacteria growing on microplastics (Jacquin *et al*., 2019). There are reports of contamination in every level of the food chain, including human beings (Cox *et al*., 2019; Monteiro *et al*., 2018; Wright & Kelly, 2017; Zhou *et al*., 2020; Zimmermann *et al*., 2021).

The scientific community has established an eco-friendly solution for some plastic types and called them biodegradable plastics; however, they have insignificant contributions in current global plastic production (Bhagwat *et al*., 2020; Borrelle *et al*., 2020; Rosenboom *et al*., 2022; Shieh *et al*., 2020; Weckhuysen, 2020). The earliest reports of biodegradation of plastic material are from the mid-1970s (Chamas *et al*., 2020; Urbanek *et al*., 2018). The various reports of biodegradable microbes are from dumping ground, open ocean, seashore, and leftovers from oil refineries due to their property to provide an optimum condition for the microbe to grow and evolve against the high concentration of plastic waste (Amobonye *et al*., 2021; Emadian *et al*., 2017; Lebreton *et al*., 2018; Zhao *et al*., 2021). Recent reports of mealworm-based plastic assimilation and the role of gut microbiota are pointing toward a large-scale worm-based solution of plastic waste (Yang *et al*., 2020; Kundungal *et al*., 2019; Kim *et al*., 2020; Bilal *et al*., 2021). On the other hand, the known biocatalysts that degrade similar linker bonds in biopolymers such as amidase, hydrolase, and esterase were also found to be effective degraders of C-X group’s plastic types. After the genomic revolution in biological sciences, metagenomic studies have changed the pipeline from microbe discovery to genome mining and further enzyme discovery and engineering (Kobras *et al*., 2021). This research work explored the feasibility of a functional bio-based solution for accumulated mixed plastic waste. The collected utilization of mixed plastic types is a prerequisite for any large-scale affordable solution (Ballerstedt *et al*., 2021). In the present manuscript, we have collected the previous reports of biodegradation of plastic material and analysed the pattern of biodegradation of different plastic types to test if a joint consortium of microbes is practically possible to degrade mixed plastic waste. Our finding has pointed towards a class coordinated behaviour of plastic types. A detailed description of the results and methodologies is provided in the manuscript’s respective sections.

## Material and Methodology

We started this work with the collection of the latest reviews on the related topics of biodegradation of plastic material, including both generic and focus on single plastic-type (Ali *et al*., 2021; Roager & Sonnenschein, 2019). The global covid-19 pandemic followed by the strict lockdown worldwide could be factors behind the sudden surge in reviews. These reviews covered a broad spectrum of enzymatic, microbial, metagenomic, and consortium development studies on various plastic types. We have prepared a database which include these individual studies reported in excellent recent reviews along with other studies reported in literature (Gambarini *et al*., 2021; Gan & Zhang, 2019), and database existing resources of collected literature (Gan & Zhang, 2019; Gambarini *et al*., 2021) are also included in the database. The latest reports not covered in the previous collection are searched *via* manual citation walking from existing reports on Google Scholar. The literature in the English language is only considered while searching. After the initial validation of relevancy of the reports, we manually mined the information about *digital object identifier* (***DOI***), the article’s topic, journal’s name, abstract, microorganism, plastic-type, microorganisms, and the enzymes from the individual research articles collected in our in-house database. The IDs of gene, protein *etc*. from the different biological databases such as Genebank (Benson *et al*., 2013), PDB (Burley *et al*., 2021), and Uniprot (UniProt Consortium, 2021) are converted into UniProt identifiers to maintain uniformity and remove the redundancy in the collected data. The name of the microorganisms is converted into NCBI taxonomical ids using the NCBI Taxa module of package ete3 (Huerta-Cepas *et al*., 2016). The taxonomical lineage of individual taxa ids is automatically fetched from NCBI using the NCBI Taxa module through a python script.

We performed two kinds of analysis, hierarchical clustering and non-parametric multidimensional scaling (NPMDS), on three different aspects of information (microbes, plastic type and enzymes etc.) available for plastic types. The open-source libraries NumPy (Harris *et al*., 2020) and pandas (https://zenodo.org/record/5893288#.YhylWpNBy3J) are utilized for the data pre-processing, scipy (Virtanen *et al*., 2020), and sklearn (https://jmlr.csail.mit.edu/papers/v12/pedregosa11a.html) are used to implement NPMDS and hierarchical clustering analysis. The jupyter notebook of the distinct analyses is attached in the supplementary material. We have information about the names of microorganisms and the sequence of enzymes known to degrade various plastic types. The information about the co-occurrence of different plastic types was gathered from in-house reports. The information is converted into association matrixes, and NPMDS and hierarchical clustering are performed to get the pattern of degradation of various plastic types. The microbial plastic degradation analysis is performed at the level of Genus to merge the distinct species-level information of individual reports. A Common hierarchical evolutionary tree for a set of taxa ids is generated from the NCBI tree browser (Schoch *et al*., 2020) to assign Genus’s relative position in the heatmaps. The limited availability of data related to number of enzymes reported to degrade is reported for different type of plastic types is non-uniform and low compared to microorganisms, leading to a sparse association matrix. A sequence-similarity linkage principle is applied to enrich the sparse matrix on different similarity thresholds. The visualization plots are developed using Matplotlib (Hunter, 2007), Seaborn (Waskom, 2021), Plotly (https://plot.ly) open source python libraries. The in-house database has converted into an interactive portal on *plasticbiodegradation*.*com* for the appropriate users to explore. The interactive interface of our database will be helpful for anyone to start literature mining on any aspect of microbial degradation of plastic materials.

## Results

### Microbial degradation result

The microbial degradation of plastic wastes in an open environment follows the process of biodeterioration, depolymerization, and assimilation. Biodeterioration of plastic material is the initial process to dismantle the polymeric surface through extracellular polymeric substances to provide the optimum condition for anchoring microorganisms. Microbes use the extracellular release of internal metabolites to change the physicochemical condition, such as surface pH, to obtain deterioration (Sharma *et al*., 2017). The non-biotic forces of the environment, such as sunlight, water, and air, also assist in priming the initial conditions. Once the microbe establishes the biofilm on the plastic surface, they release extracellular enzymes to depolymerize the plastic polymer. The depolymerization process converts more extended polymer into oligomers of a few repetitive units. The resultant oligomers are further internalized and assimilated in the metabolic process of colonizing microbes (Ghosh et al., 2019). Out of all 481 reports of microbial degradation in the in-house database, 11 reports have no information about the name of microorganisms therefore we excluded these studies from our analysis. Further, we excluded the reported studies related to biodegradation of plastic by gut microbiota, uncultured bacteria, and uncultured fungi. Out of the remaining 466 reports, 325, 157, and 11 reports contain bacteria, fungi, and metazoa species, respectively. The 27 reports have both bacteria and fungi species which were also included in study. The ubiquitous presence of bacterial degradation studies could be due to robust survival capacity of bacterial species and easy experimental handling of bacterial based biodegradation.

We have analyzed the taxonomical data of microbes at the Genus level to circumvent the differences in various studies at their taxon level identification. In our in-house data we have 218 unique genera for 80 different plastic types, and only 62 genera are reported for more than three plastic types. We only considered those plastic types reported to be degraded by more than ten unique genera, which resulted in 20 plastic types which we considered for our analysis out of 80 different types of plastic. The PLA/PHB blend was dropped from further analysis due to mixed composition. An association matrix of size (207×19) was filled with the binary values of 1 and 0 based on genus plastic types occurrence in metadata. The association matrix is further utilized to get a co-occurrence matrix of size (19×19) for all plastic-type. The Figure 2 shows the non-metric multidimensional scaling (NMMDS) and hierarchical clustering plots of selected plastic types with precomputed cosine similarity of co-occurrence matrix. The selected plastic types are clustered based on their membership of C-C or C-X type of plastic substrates. The results presented in Figure 2 indicate that PP, LDPE, HDPE, PE, and PS falls in one cluster which appears to be specific to C-C type of plastics, whereas PHB, PHBV, PLA, PES, PET, PCL, PBS, PBSA are forming another cluster of C-X type plastic material. The plastic type PU, PHA, and PHBH didn’t fall in any cluster, therefore, behave as outliers, while Nylon appears to be closer to C-C type of plastic in their biodegradation behaviour.

**Figure 1:**
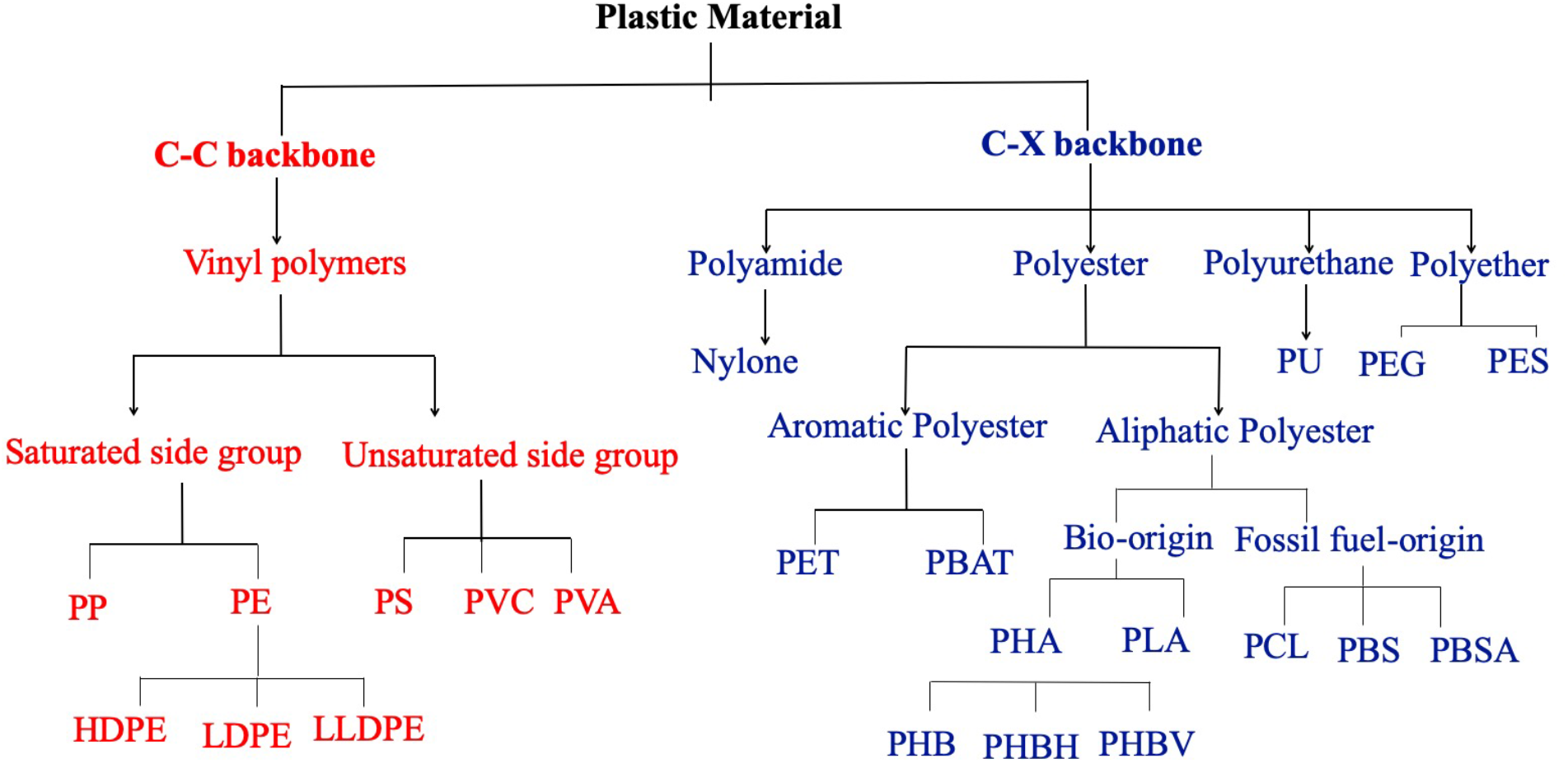
The hierarchical classification of plastic types based on atomic composition of the main backbone.

**Figure 2.**
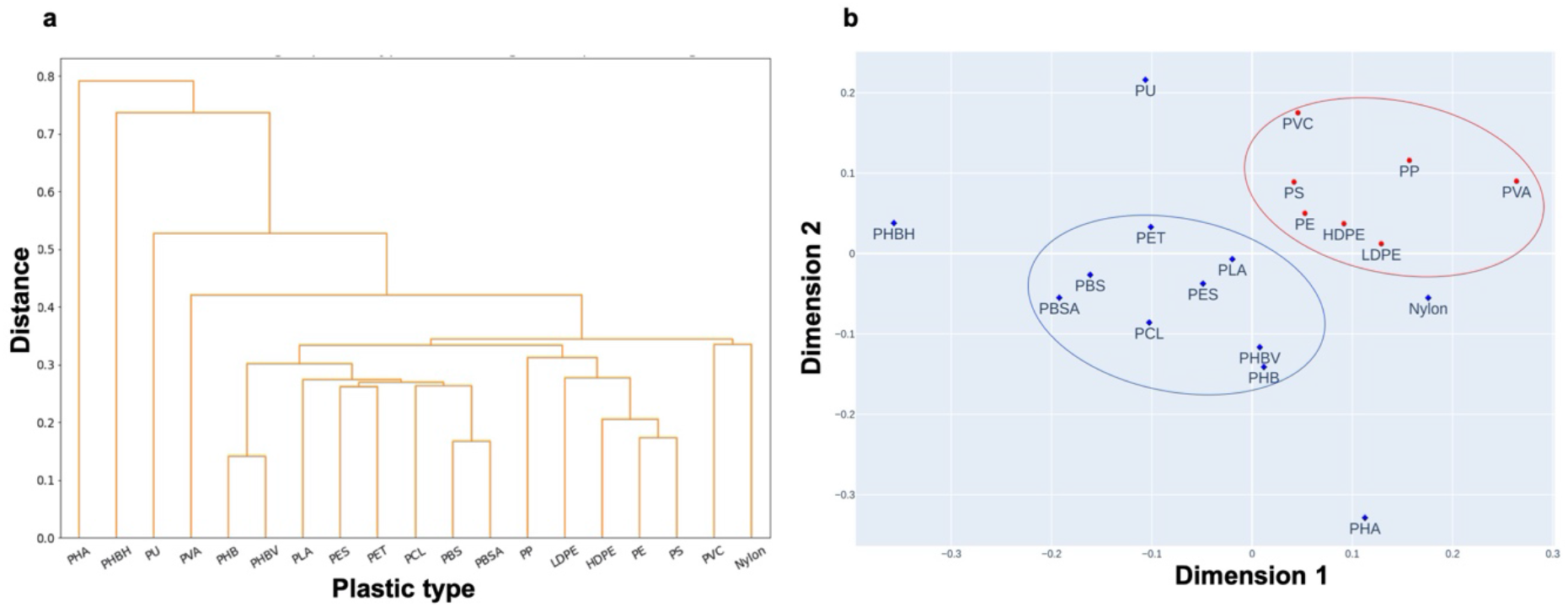
a) Hierarchical clustering and b) Non-metric multidimensional scaling (**NMDS**) plot of cosine similarity matrix computed from genus plastic association matrix where blue diamond and red circle represent C-X and C-C types of plastics respectively.

To decode the preference of bacterial genus for plastic types, we did enrichment analysis. For the same, we have selected 48 genera of bacterial origin which were reported for more than two types of above-mentioned selected plastics. The results are presented in Figure 3 as a heatmap of bacterial genus and plastic types. The colour intensity in the heatmap normalized on total entries of the column for individual plastic-type. The results indicate that the microbial degradation of plastic is not concentrated to a few specific genera but spread all over in the plot. The careful observation of the heatmap demonstrates that genus of Betaproteobacteria class (*Roseateles* to *Achromobacter*) are enriched for C-X class of plastic types, specifically PHAs. Alphaproteobacteria (*Brevundimonas* and *Rhodopseudomonas*) and Gammaproteobacteria class (*Alcanivorax* to *shewanella*) are enriched for C-C class of plastic material except *Psuedomonas* which appears to be selective for both C-C and C-X mimicking *Bacilli* class; the genus of *Bacilli* class (*Laceyella* to *Staphylococcus*) are active for both the C-C and C-X type plastic material. Overall, the genus of *Actinomycetia* class (*Thermobifida* to *Micrococcus*) is less enriched, however, appears to be selective for the C-X type plastic materials expect *Rhodococcus*. and *The Streptomyces* appears to be selective for multiple C-X type of plastic materials.

**Figure 3:**
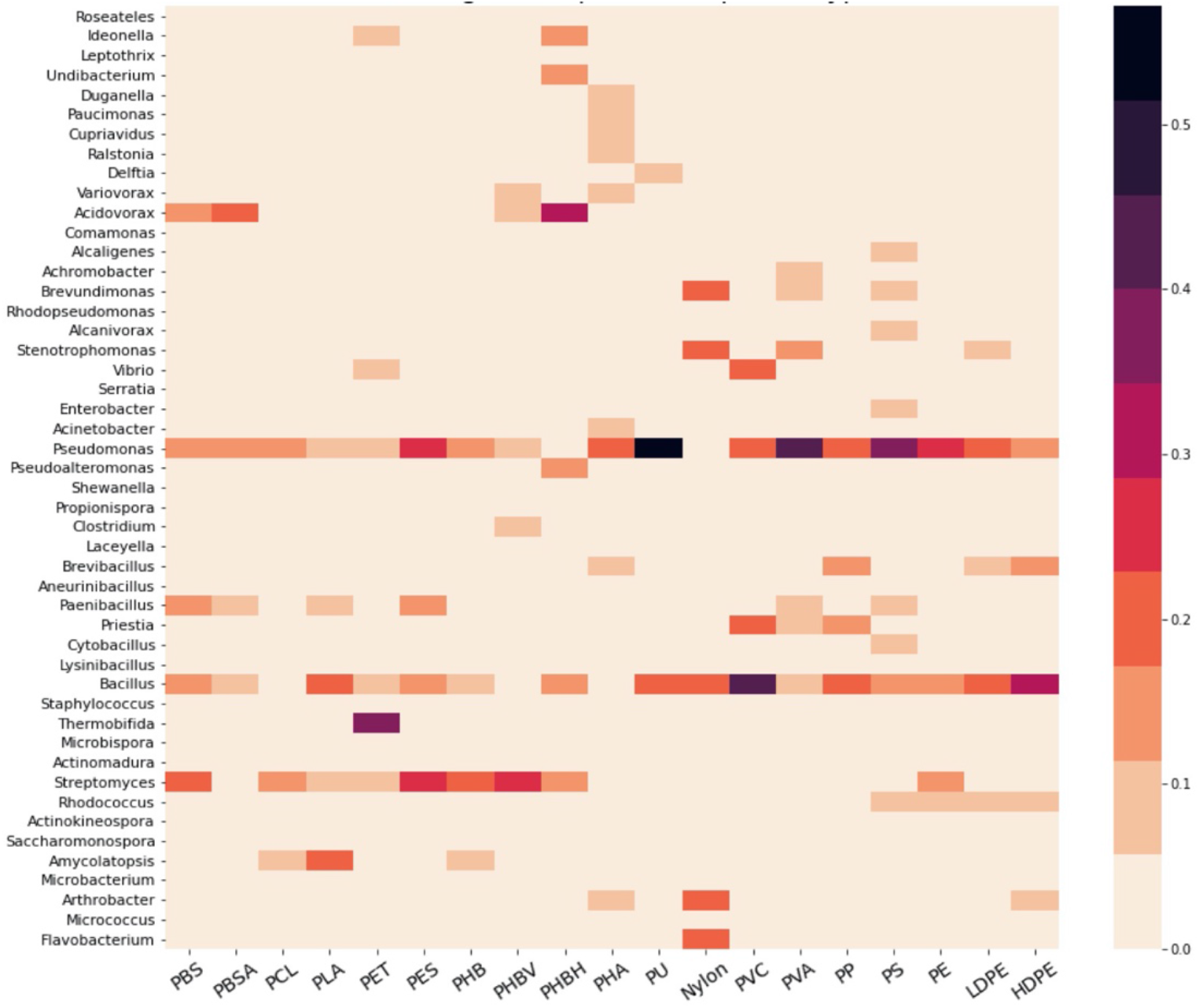
The heatmap showing the proportion of biodegradation reports of selected plastic types for different bacterial genus. The order of plastic types is arranged based on clustering analysis.

Further, to decode preferential selectivity of various fungal genus for diverse sets of plastic material, we performed enrichment analysis. Of all 79 genera of fungi kingdom associated with plastic degradation; only 28 are reported for more than two plastic-type. Figure S1 shows the heatmap of fungal genus and plastic types. We selected 42 genera of fungal origin reported for more than one selected plastic-type. Although, PHBH has no degradation report from the fungi genera therefore it is not mentioned in Figure S1. Overall the available microbial biodegradation data indicates that a smaller number of fungal genus are reported which can degrade plastics compared to bacterial genus. The data presented in FigureS1, indicates that genus of the class *Eurotiomycetes* has been extensively reported for plastic degradation; among them, the genus *Aspergillus* and *Penicillium* appears to degrade a variety of plastic materials across C-C and C-X category. According to heatmap presented in Figure S1, *Phanerodontia* genus appears to be enriched for the unsaturated C-C type plastics. The *Fusarium* genus also shows degradation capability for multiple plastic types, specifically the aliphatic ester group containing C-X class of plastic materials.

### Enzymatic degradation result

A deep understanding of microbial biocatalysts and cellular processes is necessary for developing a large-scale biotech-based solution for management of plastic waste (Hauer, 2020; Fenner *et al*., 2021; Inderthal *et al*., 2021) The continued research along this direction lead to\ reporting of many microbial biocatalysts capable of degrading plastic materials along with their microbial origin, biological pathways and sequences (Gewert *et al*., 2015; Emadian *et al*., 2017; Wei & Zimmermann, 2017). This information has been routinely being utilized for genome mining against potential microbial degraders (Viljakainen & Hug, 2021). The available information about enzymatic families are helpful to narrow down enzyme mining efforts in metagenomics studies (Danso *et al*., 2018). Rapid advancement in protein engineering methodologies, bioinformatics tool along with accessibility of diverse biological databases enables the optimization of discovered enzymes for desired function in industrial condition. Further, the discovered enzyme can be optimized for desired function (Singh *et al*., 2021). Our in-house database includes 153 enzymes which were reported previously to degrade 22 types of plastic. Our analysis indicates that the association matrix of size (153×22) is sparsely filled and contains only 7.78% of positive entries; meanwhile 70% of enzymes in association matrix are reported against only a single plastic-type. Figure S2 shows the number of enzymes for different plastic types in descending order and the distribution is observed to be skewed towards C-X type of plastics. The results indicate that highest numbers of enzymes are reported for PCL, PET, and PHB type plastic.

The enzyme’s family of cutinases, lipases, proteases, and esterases are repurposed from original biopolymer degradation to corresponding bonds containing C-X class of plastic types (Carniel *et al*., 2021; Satti & Shah, 2020). The PE, PS, LDPE, and PVA are only C-C class of plastic types in the local enzyme database associated with less than 20% of overall unique ids in the database. It appears that the oxidation of the inert C-C backbone of these plastic types, either enzymatically or physicochemically, is a must before degrading (Inderthal *et al*., 2021). The enzyme’s family of laccases, peroxidase, and hydroxylases oxidize the C-C backbone and are repurposed to degrade the C-C class of plastic types (Mitrano & Wohlleben, 2020). The non-hydrolyzable plastic types (C-C type) require consecutive action from multiple enzymes for sufficient degradation, so a whole cell-based treatment is more successful than a purified enzyme (Álvarez-Barragán *et al*., 2016). This could possibly explain the higher number of microbial biodegradation reports compared to fewer reported studies focused on enzyme based plastic degradation (Inderthal *et al*., 2021).

To enrich the sparse enzyme-plastic association matrix, we used the sequence similarity linkage to assign a plastic-type to a new enzyme with a higher E-value for two or more already known enzymes for that plastic type. The local all *vs*. all blast method has been performed on 153 enzymes, and it added 50 new connections between enzymes and plastic types at the sequence similarity threshold of e value e^-150^. Figure 4 shows the NMMDS and hierarchal clustering plots of enzyme-plastic association matrix with precomputed cosine similarity. The overall pattern of enzymatic degradation of plastic types reiterates the earlier classification and results appears to segregate the enzymes into two bigger groups specific for C-C and C-X backbone classes. The hierarchical clustering and multi-dimentional scattering data presented in Figure 4 demonstrate that C-X class is broadly divided into two sub-groups, one dominated by PHAs and the other has the rest of C-X type of plastics. Enzyme degrading PHAs also degrade PES showing the similarities in catalytic property, therefore, PES in an outgroup member in PHAs dominated group. The PU and Nylon both follow their outgroup behaviour and come close to the C-C class of plastic types which is as per our microbial degradation data reported in Figure 2. The PEG is within the C-X group of plastic types; however, it shows the distance from other polyester backbone types.

**Figure 4.**
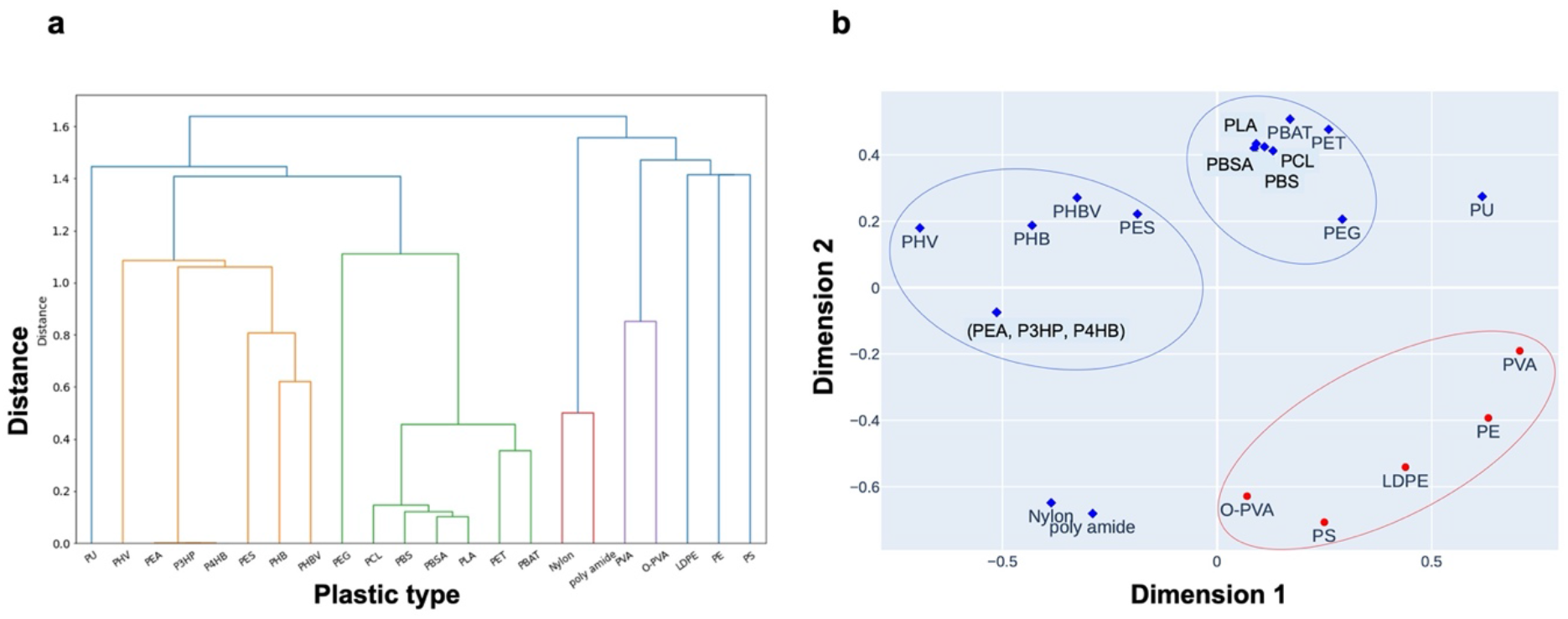
(a) Hierarchical clustering, and b) non-metric multidimensional scaling (**NMDS**) plot of precomputed cosine similarity matrix from plastic types co-occurrence data of enzyme plastic association matrix at sequence similarity threshold e^-150^ where the blue diamond and red circle symbols present the degradation profile of C-X type and C-C type of plastics respectively.

### Co-occurrence result

The co-occurrence analysis of the group of entities is a standard technique to deduce the relationship between them in the field of natural language processing (NLP) (Rozmus, 2009). We perform the co-occurrence analysis on our in-house plastic biodegradation data to find the pattern of similarity between different plastic types. There are 481 reports for 81 different plastic types; out of which 71% of studies report biodegradation of single plastic type. The Figures S3 and S4 show frequency distribution of the plastic types reported in previous studies. For co-occurrence analysis, we could select 21different plastic-type which were reported for more than five studies. A publication and plastic-type association matrix of size (446×21) is filled with binary entries based on their occurrence in the in-house database record. Later, the association matrix is used to calculate a cosine similarity matrix of size (21×21) for all selected plastic types. Figure 5 shows the NMMDS and hierarchal clustering plots of co-occurrence of different plastic types with precomputed cosine similarity. The data clearly indicates two bigger clusters corresponding to two main plastic-type category (C-C and C-X type plastics). The C-C group of plastic includes PE, PS, PP, LLDPE, HDPE, and LDPE; however, PVC and PVA show outgroup behaviour. The C-X group include PHO, PHB, PHBV, PLA, PBS, PBSA, PCL, and PES plastic types while nylon, PU, and PEG behave as outgroups compared to the rest of the C-X members. Although PET and PBAT being aromatic polyester display closeness, their similarity is not substantial. The overall co-occurrence pattern mimics the microbial and enzymatic degradation analysis patterns and supports the earlier classification of plastic types (C-X and C-C type of plastic).

**Figure 5.**
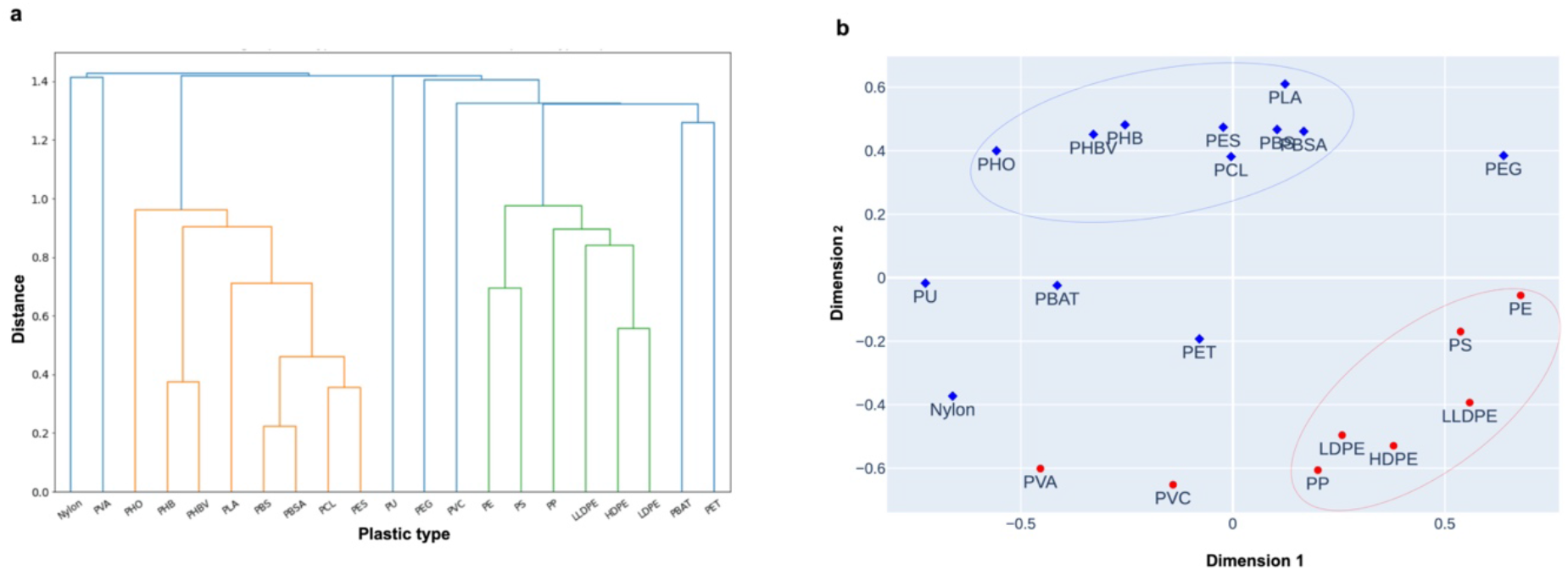
a) Hierarchical clustering, and b) non-metric multidimensional scaling (**NMDS**) plot of precomputed cosine similarity matrix from plastic types co-occurrence data of plastic publication association matrix, where the blue diamond and red circle symbols present the degradation profile of C-X type and C-C type of plastics respectively.

## Discussion

Plastic materials have been proved to be an outstanding discovery of modern time; nevertheless, their waste continuously stacks up in the open environment and turns into an ecological catastrophe. Currently, we have a limited understanding of their natural assimilation due to their extended durability and resistance against naturally eroding forces (Wilkes & Aristilde, 2017). The worldwide economies are thriving on cheap production costs and vibrant applications of plastic material (Geyer *et al*., 2017). Simultaneously, plastic waste management is underdeveloped, especially in developing countries; thus, they are becoming significant contributors of plastic waste in the open environment (Jambeck *et al*., 2015). Trashyard and incineration are the leading end-fates of used plastic with a small proportion of recycling (https://www.mckinsey.com/industries/chemicals/our-insights/how-plastics-waste-recyclingcould-transform-the-chemical-industry).

Recycling used plastic is a potential alternative to virgin plastic; however, producers are not interested in recycling every plastic type due to functional and economic factors (Garcia & Robertson, 2017; Robaina *et al*., 2020). Metropolitan waste has various kinds of plastic material with other non-plastic components; therefore, their sorting is not straightforward to implement (Lange, 2021). A pre-segregation of plastic waste in different categories is needed before the recycling process, as only a pure homologous influx is preferred over mixed waste. It is a challenging and significant bottleneck in designing a cyclic and eco-friendly solution. Using mixed plastic waste for downstream processing could be the best cost-effective move to develop an inexpensive solution (Ballerstedt *et al*., 2021).

Multiple metagenomics studies have reported a distinct microbial community called Plastisphere associated with plastic in an open environment (Kirstein *et al*., 2019). Many studies have shown that microbial strains could grow on pure plastic as their sole carbon source; however, mixed plastics are not yet thoroughly evident. We have compiled and analysed these reports to test the feasibility of a microbial degradation solution for mixed plastic waste. We first tested the similarity of microbial degradation behaviour on different plastic-type *via* different analytical methods. To get a global pattern, we further divide the plastic types into hydrolyzable(C-X) and non-hydrolyzable (C-C) groups. The results of all analysis shows that the pattern of microbial degradation follows the class segregation of plastic-type. Plastics with hydrolyzable and nonhydrolyzable backbones behave differently but are similar within the group with few exceptions such as PU, Nylon, and PET. Figure S5 shows the annual frequency of reports based on classification into hydrolyzable(C-X) and non-hydrolyzable (C-C) groups. It reveals a historic inclination towards the C-X (hydrolyzable) group of plastic-type. The pre-treatment of plastic material with physical or chemical methods can influence precise microbial degradation behaviour and, therefore, weaken the clustering patterns and lower the statistical power to determine the clusters in data. Meanwhile, lack of proper treatment reporting in all publications is the reason that we did not consider the pre-treatment factor in our analysis.

We analysed the taxonomical data of microbial strains at the genus level due to incomplete information for species level, although strains from the same Genus can vary on genomic content based on their collections site. Genus *Aspergillus, Pseudomonas*, and *Bacillus* would be perfect candidates to explore a solution for mixed plastic waste. *Rhodococcus* genus is enriched only for nonhydrolyzable backbone plastic types that would be an excellent contender for further exploration as they have a majority of stakes in overall production. The higher diversity at the genus level restricts the interpretability of our findings. This ambiguity at the genus level can only indicate the potential solutions, but search space is not limited to those genera. There are more enzymatic degradation reports for hydrolyzable plastic types than nonhydrolyzable C-C backbone types as later requires action from a cocktail of enzymes for degradation. The enzymatic degradation patterns of PHAs are different from other C-X backbone plastic types. The overall analysis showed that whole cell-based treatments are more successful than enzyme-based treatments for nonhydrolyzable C-C backbone types plastic.

The results of co-occurrence analysis are similar to microbial and enzymatic degradation analysis. The PEG, PBAT, PET, Nylon, PU, and PVA show outgroup behaviour identical to previous results. It appears that these plastic types represent an intermediatory group in between both C-C and C-X groups based on their biodegradation behaviour. Most of the reports are about the single plastic type, which led to a sparse co-occurrence matrix. Future studies with multiple plastic types are strongly recommended to have higher possibilities to find a generic solution for mixed plastic waste. Success in finding a microbial strain that is simultaneously effective on the whole plastic class, either C-C or C-X, would be taken further to a biotechnology-based solution of mixed plastic waste. Nevertheless, a consortium of microbial strains is a more feasible solution for an open environment degradation of the mixed plastic-type. The bioremediation based solutions are successfully applied in the case of a marine oil spill and heavy metal contamination (Abatenh *et al*., 2017). A similar solution would be suitable for high concentration zones of plastic waste, such as landfills and metropolis drainages. A practical and inexpensive solution to plastic waste is required for a sustainable future.

## Conclusion

Plastic waste has long-lasting tolerance against nature’s physicochemical and biological forces. The unbridled production of plastic materials combined with the irresponsible behaviour of the end-user has accelerated the harmful impacts of plastic materials in the ecosystem. A collective effort is required at different aspects of plastic material from composition to disposal to prevent this catastrophe. The scientific community has been searching for a bio-based solution for plastic waste. There are multiple reports of plastic microbial degradation, specifically from highly contaminated regions such as landfills. Based on these reports, the big corporations promote some types of plastics types as biodegradable to influence consumer’s behaviour; however, this is not entirely true. Large-scale eco-friendly, efficient methods have not yet been discovered to tackle the huge accumulation of used plastic waste. In this research work, We have done a meta-analysis on the reports of microbial degradation of various plastics. The cluster analysis of microbial degradation, enzymatic degradation, and co-occurrence of various plastic types strengthens plastic’s earlier classification into hydrolyzable(C-X) and non-hydrolyzable (C-C) groups. We observed that the number of reports from bacterial degradation is more than the fungal, and within the kingdom, the distribution is more concentrated on particular genera. The metadata genus enrichment analysis points towards a few putative genera for biodegradation solutions for mixed plastic waste. The meta-analysis indicated that a whole cell-based solution is more suitable for C-C groups as their enzymatic reports are significantly less than the C-X group of plastic types. Most of the reports contain fewer than five plastic types, which hinders the chances of finding a candidate through any metaanalysis which are capable of degrading multiple types simultaneously. The molecular mining of such strains will give insight into further engineering to develop large-scale solutions for mixed plastic. There are large discrepancies in reporting the identity of microbial strains, plastic pretreatment, and overall protocol in quantifying biodegradation in these studies. The scientific community needs to develop a standard protocol for reporting plastic biodegradation findings. An established protocol would assist in comparing and utilizing previous findings for future studies. We have compiled all the reports into an interactive web portal at *www.plasticbiodegradation.com* for easy access, which will be regularly updated in the coming future.

## Notes

http://www.plasticbiodegradation.com/

### Competing Interest Statement

The authors have declared no competing interest.

