## Supporting information for "Computational exploration of bio-remediation solution for mixed plastic waste"

**Table 1**: List of the name of plastic with their abbreviations.

| | |
|---|---|
| PE | Polyethylene |
| PP | Polypropylene |
| PVC | Poly(vinyl chloride) |
| PS | Polystyrene |
| PET | Poly(ethylene terephthalate) |
| PU | Polyurethane |
| PLA | Polylactic acid |
| PHB | Polyhydroxybutyrate |
| PHBV | Poly(3-hydroxybutyrate-co-3-hydroxyvalerate) |
| NMMDS | Non-metric multidimensional scaling |
| LDPE | Low density polyethylene |
| HDPE | High density polyethylene |
| PES | Polyethersulfone |
| PCL | Polycaprolactone |
| PBS | Polybutylene succinate |
| PBSA | Poly(butylene succinate-co-butylene adipate) |
| PHA | Polyhydroxyalkanoates |
| PHBH | Polyhydroxybutyrate-Hexanoate |
| PVA | Poly(vinyl alcohol) |
| PEG | Polyethylene glycol |
| LLDPE | Linear low density polyethylene |
| PHO | Poly (3-hydroxyoctanoic acid) |
| PEA | Poly(ethylene adipate) |
| PHV | Poly(3-hydroxyvalerate) |
| P3HP | Poly(3-hydroxypropionic acid) |
| P4HB | Poly(4-hydroxybutyrate) |
| O-PVA | Oxidized PVA |
| P(3HB-co-3HP) | Poly(3-hydroxybutyrate-co-3- hydroxypropionate) |
| P(3HB-co-3MP) | Poly (3-hydroxybutyrate-co-3-mercaptopropionate) |
| P(3HO) | Poly(3-hydroxyoctanoate-co-3-hydroxyhexanoate |
| P(HB-HV[12%] | Poly(3-hydroxybutyric acid-co-3-hydroxyvaleric acid) |

| | |
|---|---|
| PEF | Poly(ethylene furanoate or polyethylene-2,5-furandicarboxylate) |
| PC | Polycarbonate |
| P(3HB) | poly(3-hydroxybutyric acid) |
| PBAT | Poly(butylene-adipate-co-terephthalate) |
| DOI | Digital object identifier |
| ID | Identity |
| PDB | Protein Data Bank |
| UniProt | Universal Protein Resource |
| NCBI | National Center for Biotechnology Information |
| NPMDS | Non-parametric multidimensional scaling |
| NumPy | Numerical Python. |
| Pandas | Python Data Analysis Library |
| Scipy | Scientific Python |
| pH | Potential of hydrogen |

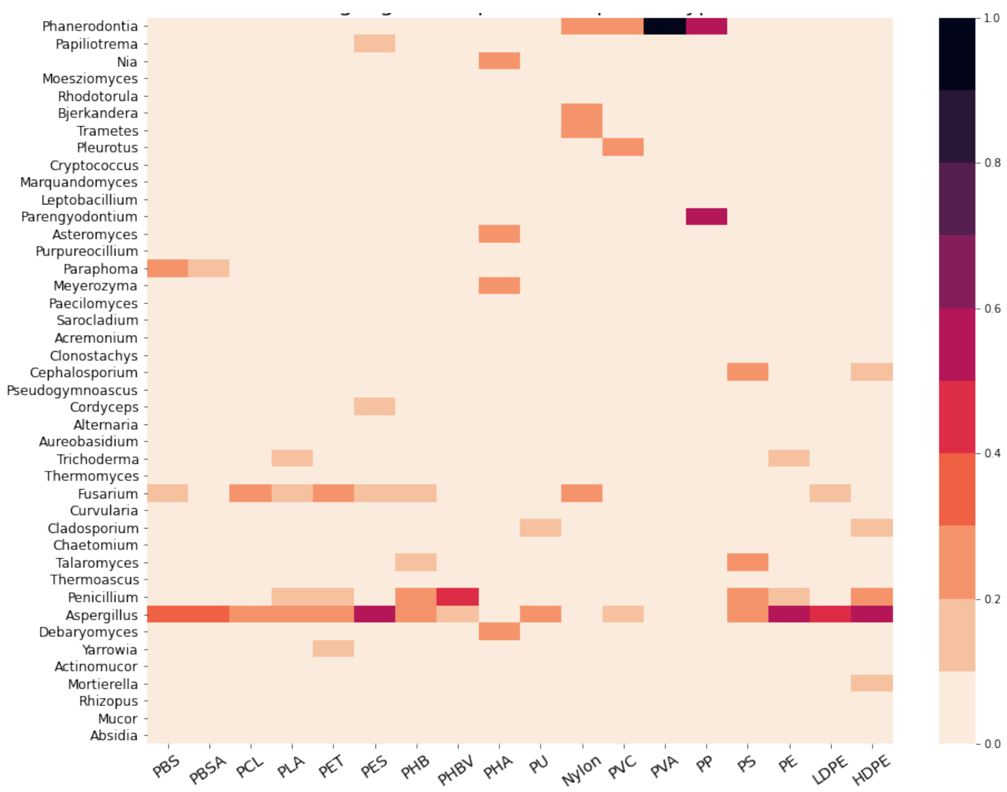


**Figure S1:** The heatmap showing the proportion of biodegradation reports of selected plastic types for the different fungal genus. The order of plastic types is arranged based on clustering analysis.

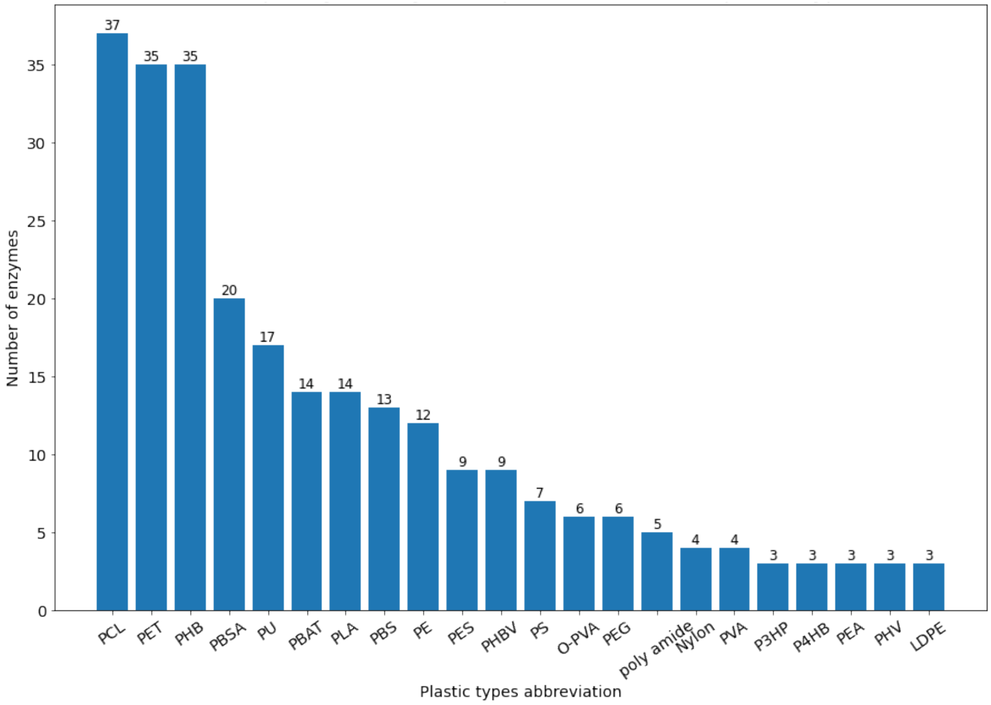


**Figure S2**. The frequency plot for different plastic types is based on the number of enzymes reported degrading corresponding plastics.

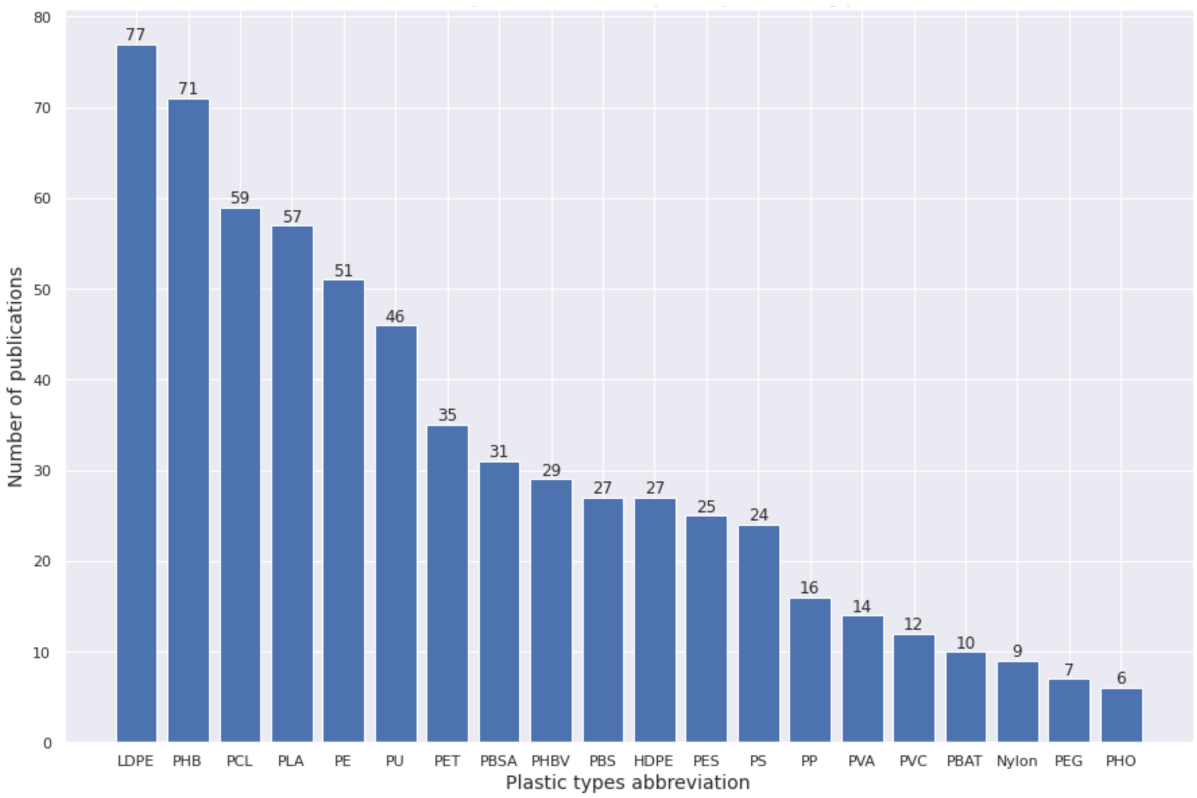


**Figure S3**. The frequency plot for different plastic types based on the number of publications that reported the biodegradation of corresponding plastics.

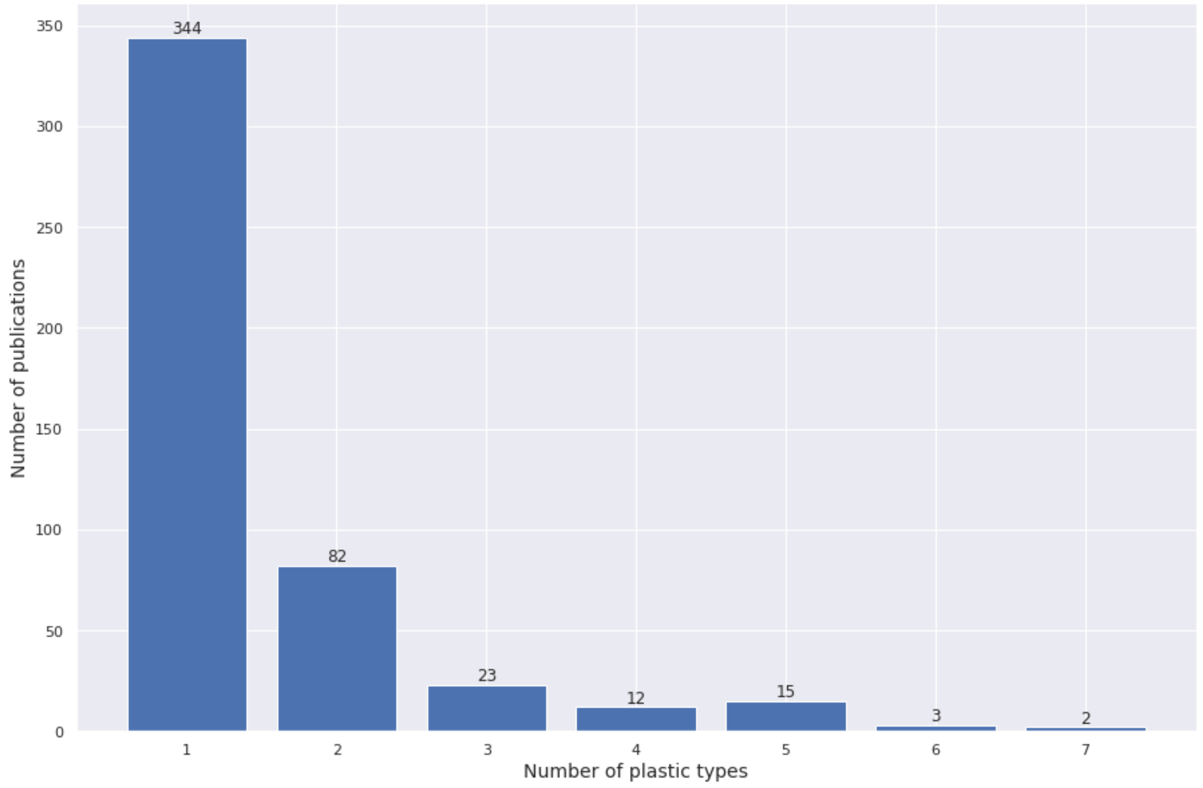


**Figure S4**. Count-plot showing the number of plastic types reported in a number of publications.

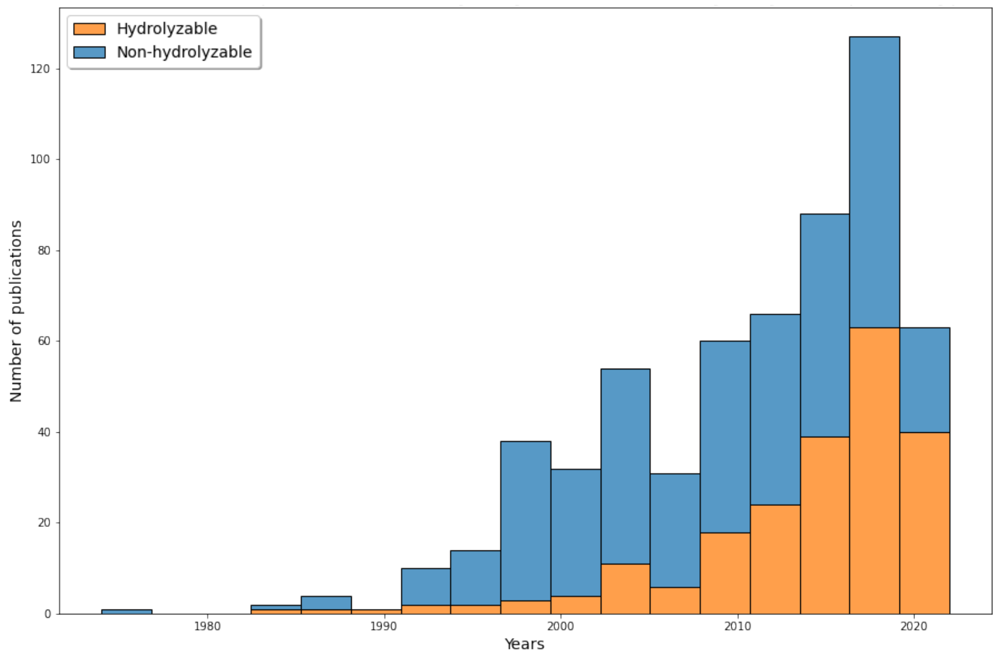


**Figure S5**. Annual trends of publication for hydrolyzable and non-hydrolyzable plastic types.
